# Short chain fatty acids potentiate azoles by reprogramming fungal acetyl-CoA metabolism

**DOI:** 10.64898/2026.06.29.735392

**Authors:** Christopher McCrory, Sofya Rabinovich, Harshini Weerasinghe, Tricia L Lo, Angavai Swaminathan, Calvin Kraupner-Taylor, Traude H Beilharz, Judith Berman, Ana Traven

**Affiliations:** Department of Biochemistry and Molecular Biology & the Infection Program, Biomedicine Discovery Institute, Monash University, Clayton 3800, VICTORIA, AUSTRALIA; Shmunis School of Biomedical and Cancer Research, George S. Wise Faculty of Life Sciences, Tel Aviv University, Ramat Aviv, ISRAEL; Department of Biochemistry and Molecular Biology & the Stem Cells and Development Program, Biomedicine Discovery Institute, Monash University, Clayton 3800, VICTORIA, AUSTRALIA

## Abstract

Pathogens colonise metabolically diverse host environments. How metabolites found in host environments regulate antimicrobial drug susceptibility remains to be fully understood. Here we report on the roles of gut metabolites, short chain fatty acids (SCFAs), in antifungal drug susceptibility of the gut commensal and fungal pathogen *Candida albicans*. A genetic screen revealed that *C. albicans* mutants in peroxisome biogenesis display increased tolerance to the antifungal drug fluconazole. Peroxisomes are important for the metabolism of SCFAs by β-oxidation, and exposure to the SCFAs butyrate and crotonate increased susceptibility and reduced tolerance to fluconazole. To understand if SCFAs inhibit fluconazole tolerance through their ability to inhibit histone deacetylases (HDACs), we compared them with the HDAC inhibitor trichostatin A. These experiments did not reveal an obvious connection between the degree of HDAC inhibition and the degree of fluconazole tolerance reduction. Exposure of *C. albicans* to crotonate and butyrate revealed transcriptional reprogramming involving remodelling of acetyl-CoA metabolism by upregulation of genes for β-oxidation, peroxisome biogenesis and intracellular transport of acetyl-CoA, while the expression of ergosterol biosynthesis genes was reduced. Since ergosterol gene expression is required to overcome fluconazole stress, these results explain how SCFAs reduce fluconazole tolerance. Taken together, our results implicate peroxisome biogenesis and metabolism in fluconazole susceptibility. We posit that balanced acetyl-CoA metabolism promotes sufficient ergosterol biosynthesis to overcome fluconazole stress and drive tolerant growth. These pathways are perturbed by metabolic changes induced by SCFAs. These findings add to our understanding of the importance of metabolic regulation in antimicrobial drug responses.

**SIGNIFICANCE:** Metabolites produced by microbiota or host cells regulate microbial metabolism, physiology and drug responses. In the gut, the human commensal and pathogen *Candida albicans* is exposed to short-chain fatty acids made by bacteria. *C. albicans* metabolises short-chain fatty acids *via* peroxisomal β-oxidation. Additionally, short-chain fatty acids change gene expression by inhibiting histone deacetylases. We found that short-chain fatty acids increase the susceptibility of *C. albicans* to the antifungal drug fluconazole and reduce fluconazole tolerance. Our mechanistic studies indicate that metabolic utilisation of short-chain fatty acids by *C. albicans* reduces fluconazole tolerance by changing acetyl-CoA metabolism and causing lower expression of ergosterol biosynthesis genes. Thus, short-chain fatty acids increase fluconazole susceptibility by changing fungal metabolism, while inhibition of histone deacetylases may play a more minor role. These findings shed light on the roles of metabolism and abundant gut metabolites in tolerance to a front-line antifungal drug.

## INTRODUCTION

Microbes colonise diverse host environments as commensal microbiota or infecting pathogens. In these environments, they encounter distinct nutrients and metabolites produced by other microbes and host cells. In turn, these metabolites affect microbial metabolic state and physiology. Moreover, metabolism affects antibiotic responses (1) and can also affect antifungal responses, although the mechanisms remain poorly understood (2).

The yeast *Candida albicans* is a great example of a microbe able to colonise diverse human body niches. It is a common commensal of the gut, oral and vaginal mucosa that can enter the bloodstream and grow in internal tissues and organs during systemic infections. Infections with *Candida* species cause just under 650,000 deaths per year in hospitalized patients (3), in addition to common infections such as vulvovaginal candidiasis in healthy individuals, with recurrent vulvovaginal candidiasis affecting an estimated 138 million women per year (4). Life-threatening *Candida* infections are treated predominantly by two drug classes: the echinocandins (β–1,3 glucan synthase inhibitors that repress cell wall synthesis) and the azoles (lanosterol demethylase inhibitors that repress ergosterol synthesis) (5). Most vulvovaginal candidiasis is treated with azoles, which are fungistatic (6).

*C. albicans* is intrinsically azole susceptible (7–9), but evolves azole resistance upon drug exposure (5, 7). Moreover, clinical isolates of *C. albicans* display both intrinsic and evolved azole tolerance characterised by trailing growth of a sub-population of cells at high drug concentrations (10). *C. albicans* can exhibit very high levels of azole tolerance (10), and in a clinical study, the level of fluconazole tolerance correlated with treatment failure and patient death (11, 12). Therefore, strategies to increase the activity of azoles by reducing resistance and/or tolerance could improve clinical outcomes.

Azole resistance is achieved by upregulating drug eflux and/or altering the drug target, lanosterol demethylase, which is encoded by *ERG11* (9). Azole tolerance remains less well understood, although it is promoted by limiting intracellular azole concentrations and activating stress response pathways that address the consequences of drug exposure, so that pathogen growth, however slow, can continue (9). For example, upregulation of sphingolipids and increased ergosterol biosynthesis overcome membrane stress caused by azoles; moreover, the chaperone Hsp90 and proteasome activity overcome azole-induced proteotoxicity, and several broad regulators of stress responses, such as calcineurin and TOR (target of rapamycin), also contribute to azole tolerance (10, 12–15), reviewed in (9).

The metabolic impacts of different host niches on *C. albicans* responses to azole drugs are not well understood. For example, in the gut, *C. albicans* is subjected to bacterial metabolites, including copious amounts of short-chain fatty acids (SCFAs), chiefly acetate, butyrate, and propionate (16). *C. albicans* metabolises SCFAs in the peroxisome *via* the β-oxidation pathway (17, 18). SCFAs have many other physiological effects including modulating gene expression by inhibiting histone deacetylases (HDACs) and altering the levels of acetylation of histones and other proteins (19). In addition to the abundant SCFAs, other SCFAs have also been studied for their ability to modulate non-acetyl acylations, such as beta-hydroxybutyrylation, crotonylation, and others (20–22). These non-acetyl modifications of histones regulate gene expression in specific ways and connect metabolic regulation with chromatin structure and transcriptional control (20, 21, 23). For example, supplementation of the SCFA crotonate increases the intracellular levels of histone crotonylation in mammalian cells, *S. cerevisiae* and *C. albicans*, and impacts on gene expression and biological responses (21, 23, 24). Several reports suggested that the SCFAs butyrate and acetate potentiate azoles, including via HDAC inhibition, but the mechanisms remain to be fully understood, and the impacts of other SCFAs have not been studied so far (25–27).

Here, we employed an *in vivo* transposon mutagenesis screen, coupled with transcriptomics and phenotypic analyses, to further elucidate the mechanisms of azole susceptibility in *C. albicans* and the role of SCFAs in fluconazole resistance and tolerance.

## RESULTS

### A transposon mutagenesis screen identifies peroxisome biogenesis genes as mediators of fluconazole susceptibility

To identify modulators of azole tolerance in *C. albicans*, we performed an *in-vivo* transposon mutagenesis screen as described in (28). The screen was performed in technical triplicates with a high fluconazole concentration of 10 µg/ml fluconazole as well as no-drug control conditions (**Figure 1A**). One fluconazole-treated sample revealed an enrichment of transposon-insertion mutants in genes required for peroxisome biogenesis (**Dataset S1**, **Figure 1B**), indicating that their inactivation promotes growth in the drug. Enriched genes encoded homologs of *S. cerevisiae PEX1*, *PEX6, PEX11, PEX8, PEX2, PEX14, PEX28* and *PEX32*, which are required for the biogenesis, morphology and size regulation of the yeast peroxisome (29). Of note, role of *PEX1* in peroxisome biogenesis in *C. albicans* has been demonstrated (30), while the putative functions of the other *PEX* genes are inferred from their *S. cerevisiae* homologs.

**Figure 1.**
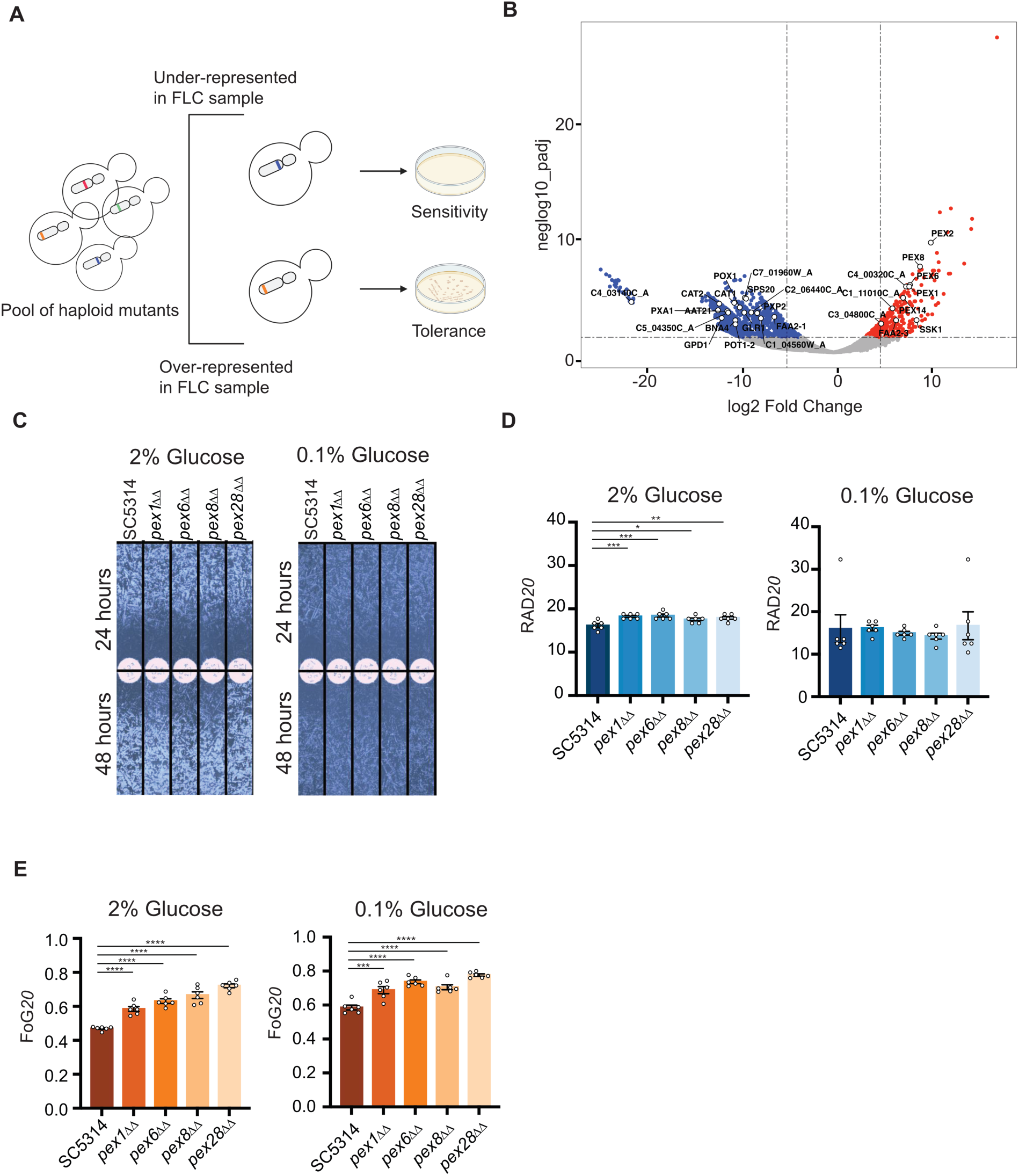
A transposon mutagenesis screen links peroxisome biogenesis to fluconazole tolerance. A. Schematic of the transposon mutagenesis screen to reveal genes involved in fluconazole susceptibility. B. Volcano plot of mutants enriched in the presence of fluconazole or control conditions. Genes related to peroxisome biogenesis and function are indicated. Also see **Dataset S1**. C. Fluconazole disk diffusion assays of *pex* mutants in synthetic complete (SC) medium with 2% or 0.1% glucose (pH5.4, buffered with HEPES) at 30 °C. D. Quantification of RAD_20_ from experiments in panel C. Shown are the average and SEM of n=6 repeats, *p<0.05, **p<0.01, ***p<0.001, ****p<0.0001 (Ordinary one-way ANOVA with Dunnett’s multiple comparisons test). E. Quantification of FoG_20_ from experiments in C. Shown are the average and SEM of n=6 repeats, ***p<0.001, ****p<0.0001 (Ordinary one-way ANOVA with Dunnett’s multiple comparisons test).

In fungi, the peroxisome is the main site of fatty acid breakdown by β-oxidation. The screen also identified a mutant in *FOX2,* the central enzyme of β-oxidation, that was enriched in fluconazole conditions, although the p-value was just outside the 0.05 cut-off (p=0.055634) (**Dataset S1**). Other genes enriched or depleted in the screen (and thus might inhibit or promote growth in fluconazole) included enzymes of β-oxidation and the glyoxylate cycle (**Dataset S1**). These results encouraged us to test the hypothesis that multiple aspects of peroxisome function are important for fluconazole responses in *C. albicans*.

We directly tested the effect on fluconazole susceptibility and tolerance with homozygous deletion mutants lacking *PEX1*, *PEX6* and *PEX8* (important for peroxisome biogenesis) or *PEX28* (important for peroxisome size, number and distribution) using disk diffusion assays. In disk assays, we measure susceptibility using RAD_20_ (i.e. radius of the zone of inhibition in mm at 20% inhibition of fungal growth), which is inversely related to resistance, and we measure tolerance as FoG_20_ (fraction of growth within the radius of inhibition relative to growth outside of it). Since glucose represses peroxisome biogenesis (29), we compared the drug responses of *pex* mutants in 2% glucose as well as 0.1% low glucose medium, which may also be more physiologically relevant (**Figure 1C-1E**). In 0.1% glucose, fluconazole tolerance increased in all strains (including the wild type) (compare **Figure 1E**), suggesting that glucose metabolism and/or signalling affect tolerance. Fluconazole resistance levels remained similar in both 2% glucose and 0.1% glucose (**Figure 1D**).

Deletion mutants lacking any one of these *PEX* genes had increased tolerance in both 2% glucose and 0.1% glucose (**Figure 1E**), consistent with the Tn screen. Interestingly, these *pex* mutants display decreased drug resistance (higher RAD_20_) relative to the wild type strain in 2% glucose, but in 0.1% pex mutant showed the same resistance as the wild type (**Figure 1C**). The uncoupling of resistance and tolerance responses in 2% glucose medium highlights the distinct mechanisms underpinning these drug responses and suggests that *PEX* genes have complex roles in fluconazole responses. Collectively, these data show that *C. albicans* peroxisome biogenesis genes modulate fluconazole tolerance and the effect is more evident in low glucose concentrations.

### SCFAs potentiate fluconazole against *C. albicans* and reduce tolerance

SCFAs are abundant gut metabolites that *C. albicans* metabolises *via* the β-oxidation pathway; thus, providing SCFAs to *C. albicans* cells is expected to increase β-oxidation and peroxisome biogenesis (17, 18). In 0.1% glucose media, acetate decreased fluconazole resistance (increased RAD_20_), while butyrate, propionate and lactate displayed a similar trend, although it was not statistically significant (**Figure 2A**). Acetate and butyrate also decreased fluconazole tolerance slightly (reduced FoG_20_), while in lactate, fluconazole tolerance increased slightly (**Figure 2A**).

**Figure 2.**
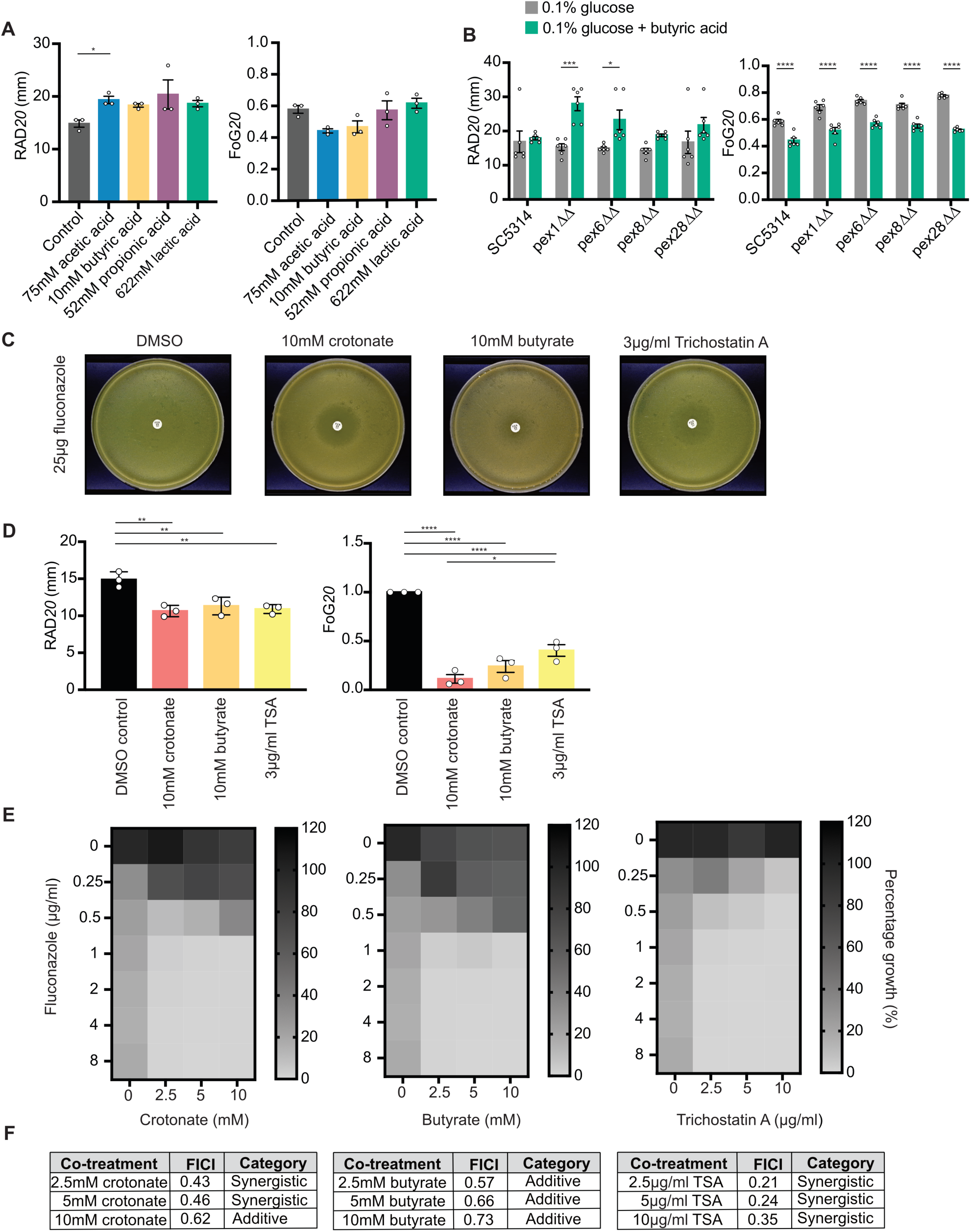
SCFAs potentiate fluconazole and inhibit drug tolerance. A. Fluconazole disk diffusion assays of *C. albicans* in SC medium (0.1% glucose, pH5.4, buffered with HEPES) at 30 °C with or without SCFAs at the indicated concentrations. Shown are averages and SEM for RAD_20_ and FoG_20_ from n=3 repeats, *p<0.05, **p<0.01, ***p<0.001 (Kruskal-Wallis with Dunnett’s multiple comparisons test). B. Fluconazole RAD_20_ and FoG_20_ for the indicated strains was determined using disk diffusion assays in SC medium with 0.1% glucose (pH5.4, buffered with HEPES) at 30 °C, +/- 10 mM butyrate. The data without butyrate is the same as in Figure 1D. All conditions were assayed in parallel but shown separately for clarity. Shown are the averages and SEM for n=6 repeats. Two-way ANOVA for RAD showed a significant effect of butyrate (p<0.0001) but no significance for genotype or interaction. Two-way ANOVA for FoG showed a significant effect of butyrate and genotype (p<0.0001) and a significant interaction (0.001). A post-hoc Sidak’s multiple comparison test was used to assess significance of butyrate treatment for each strain for RAD and FoG (*p<0.05, ***p<0.001, ****p<0.0001). C. Fluconazole disk diffusion assays in M199 medium with or without crotonate, butyrate or TSA at the indicated concentrations. DMSO was used as the vehicle control. Plates were incubated at 37 °C and photographed after 48 h. Shown are representative images from three biological repeats. Two other repeats are shown in **Figure S2**. D. Quantification of RAD_20_ and FoG_20_ from disk diffusion assays in B. Shown are the mean and SEM from three independent experiments. Only significant statistical comparisons are indicated. **p<0.01, ***p<0.001, ****<0.0001 (Ordinary one-way ANOVA with Tukey’s multiple comparisons test). E. Checkerboard assays of fluconazole in combination with crotonate, butyrate or TSA at the indicated concentrations in medium M199. The heatmaps reflect absorbance measured at OD_600nm_ at 24 h and were plotted as a percentage of growth compared to the untreated control in M199 alone supplemented with equivalent volumes of DMSO. F. Fractional inhibitory concentration index (FICI) determinations were performed as described in Materials and Methods, using the FLC MIC of 1 µg/ml. FICI values ≤0.5, 0.5 to <1, 1 to 4 or >4 are characterised as synergistic, additive, indifferent or antagonistic, respectively.

To further understand the roles of SCFAs in fluconazole responses and their possible connection to peroxisomal metabolism, we measured the effects of butyrate on fluconazole susceptibility using the *pex* mutants. Butyrate reduced tolerance and resistance in *pex* mutants, and reduced tolerance in the wild type strain (**Figure 2B**). These results suggest that butyrate potentiates fluconazole in a manner that could be both dependent and independent of peroxisomal metabolism.

Because butyrate is a strong inhibitor of histone deacetylases (HDACs) (21), it has been proposed to potentiate azoles via HDAC inhibition (26). We previously found that the SCFA crotonate, like butyrate, inhibits *C. albicans* filamentation (17), although it is only a weak HDAC inhibitor (21). We therefore reasoned that using crotonate alongside butyrate may help disentangle the roles of SCFAs in HDAC inhibition and fluconazole responses. In addition, we also included the HDAC inhibitor trichostatin A (TSA), along with crotonate and butyrate at concentrations that increase histone acetylation levels in *C. albicans* (23). The experiments were performed using medium M199 at 37 °C. M199 contains 0.1% glucose and an acidic pH of 5.4, conditions in which SCFAs have a strong effect on *C. albicans* filamentation (17). Western blot experiments confirmed that the levels of histone H3 lysine 9 acetylation (H3K9ac) increased in response to TSA or butyrate but not in response to crotonate (**Figure S1A and S1B**). At the concentrations used, the biggest increase in H3K9ac was with TSA, followed by butyarate and then crotonate with no effect (**Figure S1B**). As expected, fluconazole alone did not affect H3K9ac levels (**Figure S1A and S1B**).

In M199 medium, *C. albicans* had high fluconazole tolerance levels (FoG_20_ in disk assays) (**Figure 2C**). Crotonate, butyrate and TSA inhibited this tolerant growth and cleared the zone of inhibition (**Figure 2C and 2D, Figure S2)**. Strikingly, crotonate, the weakest HDAC inhibitor, inhibited fluconazole tolerance more than TSA, the strongest of the HDAC inhibitors, (**Figure 2C and 2D**), suggesting that HDAC inhibition cannot be the only mechanism by which SCFAs decrease fluconazole tolerance. Furthermore, a small reduction in the RAD_20_, was also evident in the disk assays in M199 medium, indicating increased resistance (**Figure 2D**). This again highlights the complex effects of SCFAs on fluconazole resistance and tolerance detected with disk diffusion assays.

To further address the effects of SCFAs and TSA, we performed checkerboard assays in liquid culture using the CLSI approach (35 °C, 24 or 48 h of growth, modified to use M199 instead of RPMI), to determine the fluconazole minimum inhibitory concentration (MIC) and any synergistic or additive effects of the compounds. Under these conditions neither crotonate, butyrate nor TSA alone reached an MIC_80_ (i.e. 80% growth inhibition) (**Figure 2E and S3A**). For fluconazole, initial response was dose-dependent growth inhibition, reaching MIC_80_ at 1 µg/ml (**Figure S3A**). In these liquid assays with M199 medium, substantial trailing growth at higher fluconazole concentrations (**Figure S3A**) was consistent with the high tolerance observed in disk diffusion assays performed with M199 plates (**Figure 2C**).

In fluconazole concentrations of 1 µg/ml and higher, the addition of SCFAs increased growth inhibition relative to fluconazole alone, with TSA inhibiting growth at all fluconazole concentrations tested (**Figure 2E and S3B-S3D**). FICI calculations identified synergistic interactions for crotonate and TSA with fluconazole, and additive interactions for butyrate with fluconazole (**Figure 2F**). Collectively, these experiments show that, in combination with fluconazole, SCFAs and TSA increase fluconazole susceptibility and reduce tolerance.

### The effects of SCFAs on fluconazole susceptibility in the hyphae-defective *cph1Δ/Δ efg1Δ*/*Δ* mutant

Filamentous growth of *C. albicans*, like tolerance, is affected by many factors. Because, like fluconazole, crotonate and butyrate inhibit *C. albicans* hyphal growth (17, 25, 31–33), we asked if their effect on filamentous growth and on fluconazole tolerance are related by analysing the hyphae-deficient *cph1Δ/Δ efg1Δ*/*Δ* mutant (34, 35). As expected, the *cph1Δ/Δ efg1Δ*/*Δ* mutant does not make hyphae in M199 medium (**Figure 3A**). In disk diffusion assays, the *cph1Δ/Δ efg1Δ*/*Δ* mutant exhibits tolerance (**Figure 3B**). Crotonate inhibited fluconazole tolerance of the *cph1Δ/Δ efg1Δ*/*Δ* mutant at 30 °C and 37 °C (**Figure 3B**). This result suggests that the effect of crotonate on hyphal growth is distinct from its effects on fluconazole tolerance.

**Figure 3.**
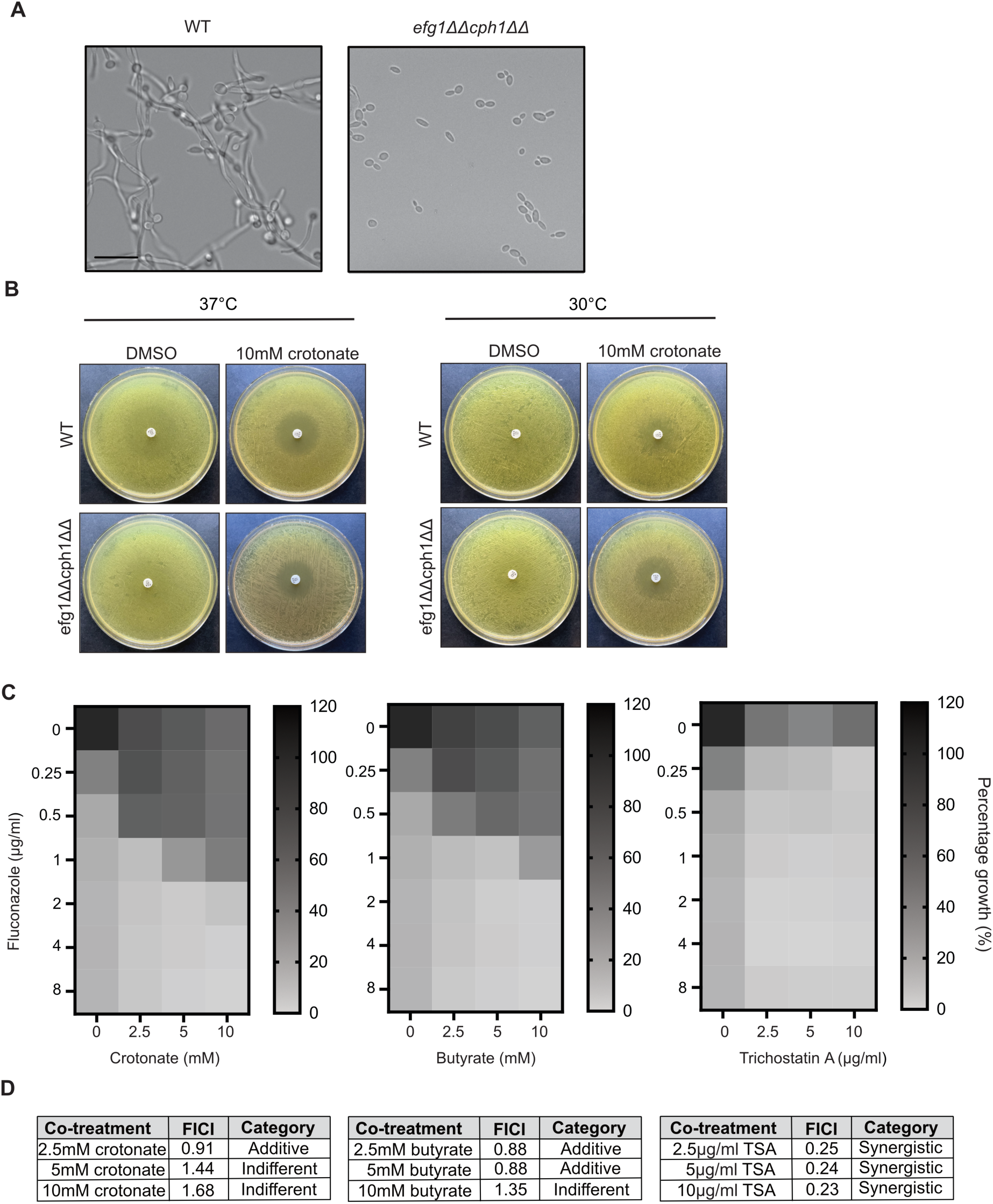
The effects of SCFAs and TSA on fluconazole susceptibility in a hyphae-defective mutant. **A.** Hyphal morphogenesis of wild type and *cph1Δ/Δ efg1Δ*/*Δ* mutant *C. albicans* in liquid M199 at 37 °C. Images were taken at 40X magnification. Scale bar is 20 μm. **B.** Fluconazole disk diffusion assays in M199 with or without 10 mM crotonate or equivalent DMSO vehicle control. Plates were incubated at 30 °C or 37 °C and photographed after 48 h of incubation. Shown are the representative images of two biological repeats. **C.** Checkerboard assays of the *cph1Δ/Δ efg1Δ*/*Δ* mutant with fluconazole in combination with crotonate, butyrate or TSA at the indicate concentrations in medium M199. Heatmaps reflect absorbance measured at OD_600nm_ at 24 h. Graphs were plotted as a growth percentage compared to the untreated control with DMSO vehicle only. **D.** Fractional inhibitory concentration index (FICI) from data in C. FICI values ≤0.5, 0.5 to <1, 1 to 4 or >4 are characterised as synergistic, additive, indifferent or antagonistic, respectively.

In liquid MIC assays, the *cph1Δ/Δ efg1Δ*/*Δ* mutant was somewhat susceptible to crotonate and butyrate, with reduced growth observed at higher SCFA concentrations (**Figure S4A**). However, the MIC_80_ was not reached. At the highest concentrations used (20 mM), crotonate and butyrate only inhibited *cph1Δ/Δ efg1Δ*/*Δ* growth by about 60% (**Figure S4A**). FICI calculations showed that TSA displayed synergistic effects with fluconazole against the *cph1Δ/Δ efg1Δ*/*Δ* mutant, while crotonate and butyrate achieved either additive or indifferent effects (**Figure 3C and 3D**).

### The transcriptional response of *C. albicans* to SCFAs and TSA

To understand the mechanism of fluconazole potentiation by SCFAs and TSA, we performed RNAseq on cells briefly exposed to fluconazole (3 µg/ml) in the presence or absence of SCFAs or TSA. We chose the 30 min exposure time to avoid indirect effects on the transcriptome due to prolonged drug stress, and no growth inhibition was detected under these conditions (**Figure S5**). The complete RNAseq dataset can be analysed interactively <u>here</u>. **Datasets S2-S5** detail the analyses of the RNAseq data.

Fluconazole samples clustered with the solvent-control (DMSO) samples in multi-dimensional scaling (MDS) plots (**Figure 4A**), consistent with the idea that the brief fluconazole exposure did not cause a major transcriptional response. Nonetheless, fluconazole did induce the expression of ergosterol biosynthesis genes (**Figure 5, Dataset S4**), indicating that the cells sensed the drug. SCFA-treated samples (butyrate and crotonate) clustered more closely together, with or without fluconazole (**Figure 4A**). Moreover, the transcriptional response to the SCFAs was larger than for the TSA-treated samples both in the presence of fluconazole (comparing SCFAs+fluconazole and TSA+fluconazole combinations to fluconazole alone samples) (**Figure 4B**), and in the absence of fluconazole (comparing SCFAs and TSA to DMSO solvent control samples) (**Figure S6A**). There was more overlap between crotonate and butyrate-regulated genes, than between the SCFAs and TSA, both in the presence and absence of fluconazole (**Figure 4C, Figure S6B, Datasets S2 and S3**). For example, in the presence of fluconazole, 456 genes were upregulated by both butyrate and crotonate, 345 genes were up-regulated by both butyrate and TSA, while 243 genes were up-regulated by both crotonate and TSA (**Figure 4C**). Similarly, 507 genes were down-regulated by both crotonate and butyrate, 301 genes were down-regulated by both butyrate and TSA, while 225 genes were down-regulated by both crotonate and TSA (**Figure 4C**). The larger overlap between genes regulated by both butyrate and TSA relative to genes regulated by both crotonate and TSA aligns with the fact that butyrate is a stronger HDAC inhibitor than crotonate. We also performed pair-wise comparisons which showed no differentially expressed genes at FDR of 0.05 and 1.5 fold change for crotonate versus crotonate + fluconazole, 11 genes for butyrate versus butyrate + fluconazole, and 240 genes in TSA versus TSA+fluconazole (see <u>here</u> and **Dataset S5**). This is consistent with distinct responses to SCFA versus TSA and further shows that, in the presence of crotonate or butyrate, the transcriptional response is largely driven by the SCFAs, rather than by the brief exposure to fluconazole.

**Figure 4.**
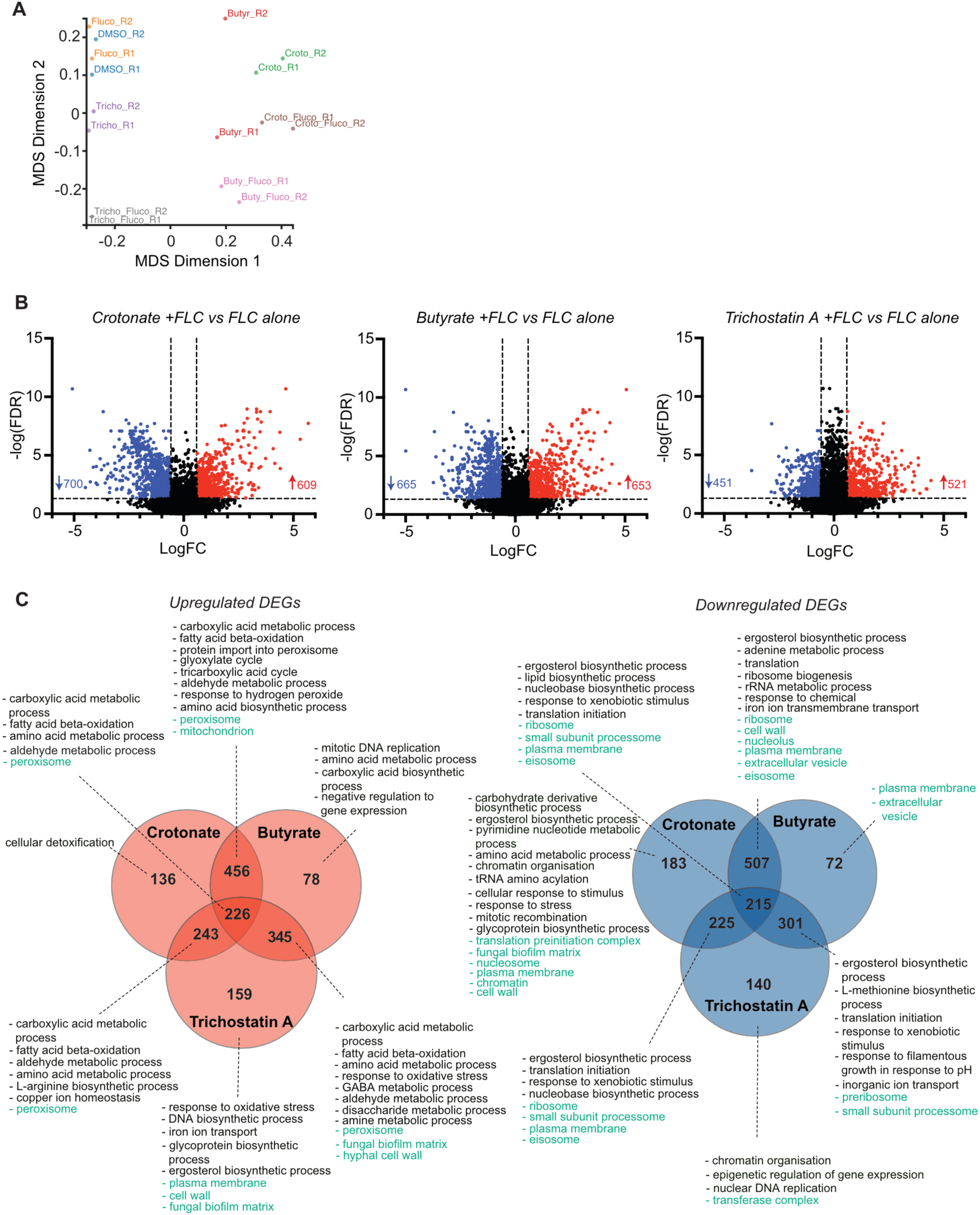
SCFAs cause a large transcriptional reprogramming in C. albicans. **A.** Multidimensional Scaling plot of the RNAseq data. **B.** Volcano plots of differentially expressed genes following treatment with crotonate, butyrate or TSA in the presence of fluconazole relative to fluconazole alone (FDR 0.05, log_2_ FC 1.5). **C.** Venn diagrams for genes differentially expressed by combinations of butyrate, crotonate and TSA with fluconazole relative to fluconazole alone. Gene Ontology (GO) Term Finder analyses was performed at the Candida Genome Database (p <0.1). Many functionally overlapping GO terms for Process (black) and Component (green) were identified. The functional groups were summarised in the figure to avoid repetition. The complete GO analyses is in **Datasets S3**.

**Figure 5.**
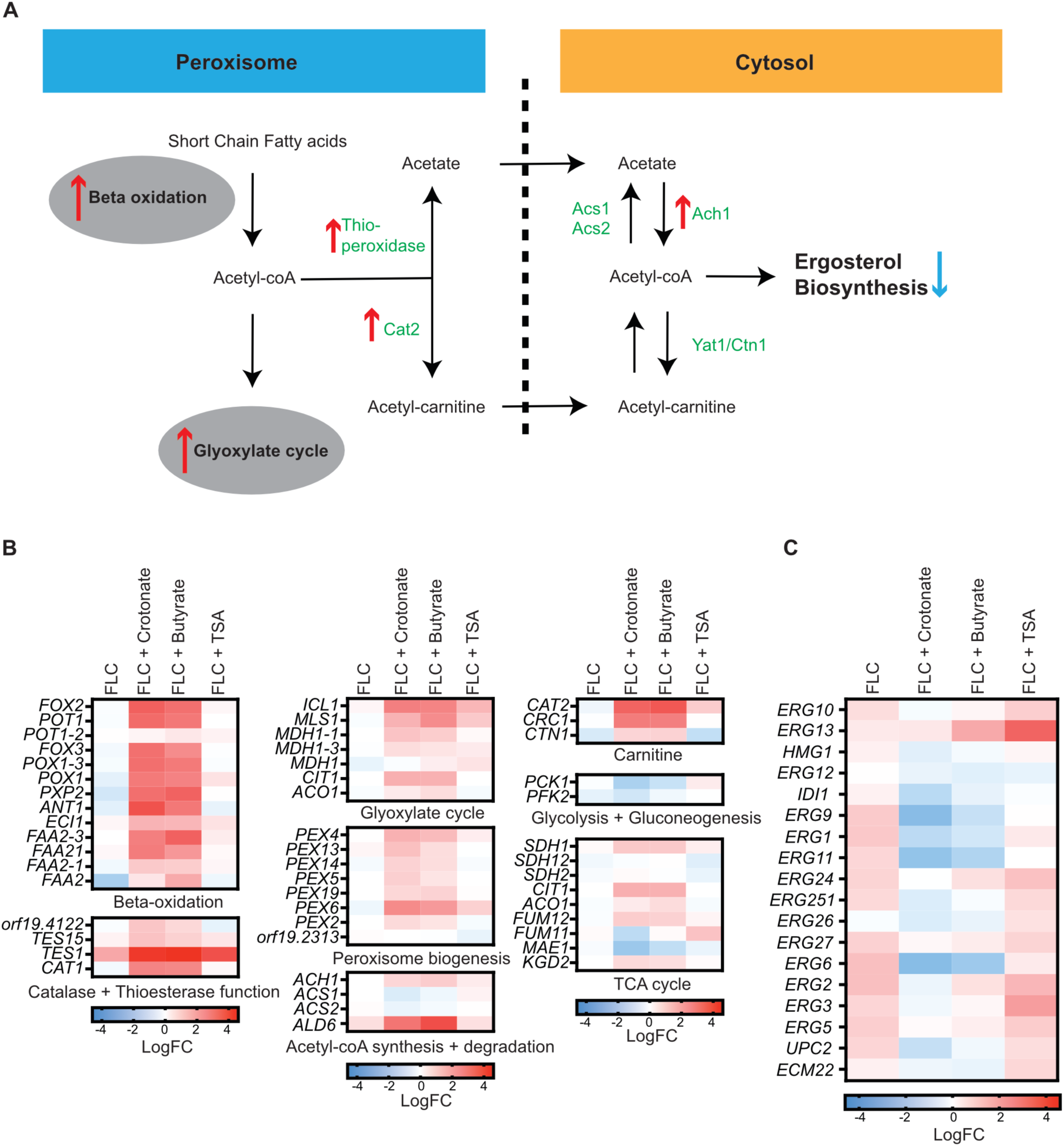
SCFAs remodel acetyl-CoA metabolism and reduce ERG gene expression. **A.** Schematic of acetyl-CoA metabolism in fungi in the peroxisomal and cytosolic compartments. The pathways transcriptionally up-regulated by the SCFAs are shown with a red arrow, while those down-regulated by SCFAs are shown with a blue arrow. Acetyl-CoA also enters the mitochondria where it is metabolised in the Krebs cycle (not depicted). **B.** Heat map of acetyl-CoA metabolism genes in the presence of fluconazole +/-crotonate, butyrate or TSA. Gene expression is expressed relative to the solvent control condition (DMSO). FLC – fluconazole; TSA – trichostatin A. Data used to create the heat map is shown in **Dataset S4**. **C.** Heat map of *ERG* gene expression in response to fluconazole (FLC), crotonate +FLC, butyrate + FLC or TSA + FLC. The data is expressed relative to the solvent control (DMSO). The false discovery rate (FDR) was 0.05. Data used to create the heat map is shown in **Dataset S4**.

For all three compounds (crotonate, butyrate or TSA) in combination with fluconazole relative to fluconazole alone genes encoding proteins involved in carboxylic acid and amino acid metabolism, peroxisomal β-oxidation, surface structures and stress responses were enriched in gene ontology (GO) terms (**Figure 4C, Dataset S3**). Genes encoding proteins involved in ergosterol biosynthesis as well as in ribosome biogenesis, translation and DNA processes were depleted, suggesting initial stages of growth inhibition (**Figure 4C, Dataset S3**). Similar functional groups were also enriched with SCFAs and TSA in the absene of fluconazole (**Figure S6**). Collectively, these expression changes are consistent with metabolic reprogramming and growth stress in *C. albicans* treated with SCFAs or TSA.

Of note, genes expressed in hyphae, such as *ECE1* and *SOD5* were down-regulated in the presence of SCFAs but less so in TSA, in the presence and absence of fluconazole (**Dataset S2, S3 and** <u>here</u>). This is consistent with hyphal inhibition by SCFAs (17, 25, 33) and distinct transcriptional effects of SCFAs versus TSA. Moreover, the SCFAs but not TSA increased the expression of the Major Facilitator Superfamily (MSF) transporters *NAG3* and *NAG4* (**Dataset S2, S3 and** <u>here</u>). The putative polyamine transporter *TPO3* was up-regulated more in SCFAs than in TSA (**Dataset S2, S3 and** <u>here</u>). **T**he *NAG3* and *NAG4* transporters modulate resistance to a number of compounds (36, 37), and *TPO3* may also be involved in eflux of certain chemicals (38). However, given that SCFAs cause increased expression of these transporters, but increase fluconazole susceptibility, it is unlikely that these transporters are efluxing azoles under these conditions. Rather, as Nag3 and Nag4 have been implicated in sugar uptake (36), we posit that *C. albicans* may use them to import SCFAs.

### SCFAs modulate acetyl-CoA metabolism and lower the expression of ergosterol biosynthesis genes

The GO analyses in **Figure 4** showed that SCFAs and TSA alter the expression of metabolic genes. Especially affected were genes involved in acetyl-CoA metabolism, including β-oxidation, peroxisome biogenesis and ergosterol biosynthesis. Since fluconazole inhibits ergosterol biosynthesis, we asked if these metabolic effects could explain how SCFAs and/or TSA increase fluconazole susceptibility by examining these transcriptional changes in more detail.

An overview of the relevant metabolic pathways is shown in **Figure 5A**. Peroxisomal β-oxidation produces acetyl-CoA, which is transported to the cytoplasm and into the mitochondria for the Krebs cycle. Peroxisomal acetyl-CoA also enters the glyoxylate cycle. Since acetyl-CoA cannot traverse membranes, it is transported from the peroxisome through the carnitine shuttle (39). In addition to the carnitine shuttle, acetyl units can also be transported from the peroxisome as acetate, produced from acetyl-CoA by peroxisomal thioesterases (40). In the cytoplasm, acetate is converted back to acetyl-CoA by acetyl-CoA synthetase. Cytoplasmic acetate is also produced by glucose fermentation. The cytoplasmic pool of acetyl-CoA is used to synthesise ergosterol, among other molecules (41).

Exposure to fluconazole alone did not alter β-oxidation or peroxisome gene expression (**Figure 5B, Dataset S4**). Exposure to SCFAs either alone or in combination with fluconazole strongly up-regulated the β-oxidation pathway, as well as the peroxisomal catalase *CAT1* and several *PEX* genes involved in peroxisome biogenesis (**Figure 5B, Dataset S3 and S2**). By contrast, TSA had only minor effects on these peroxisome-associated pathways (**Figure 5B**). Other pathways up-regulated by SCFAs, and to a lesser extent by TSA, included the glyoxylate cycle, some mitochondrial TCA cycle enzymes (**Figure 5B, Dataset S4**), as well as enzymes that catalyse the conversion of acetyl-CoA to acetyl-carnitine (*CAT2*), and thioesterases (e.g. *TES1* and *TES15*), which are thought to catalyse the peroxisomal conversion of acetyl-CoA to acetate (40) (**Figure 5B, Dataset S4**). Collectively, these data suggest that SCFA a promotes acetyl-CoA formation and the transit of converted acetyl units from the peroxisome to the cytoplasm (40).

As expected, fluconazole inhibition of ergosterol biosynthesis caused a compensatory up-regulation of *ERG* genes that encode ergosterol biosynthesis enzymes (42). This result indicates that the cells sensed and responded to fluconazole stress within the 30 min of drug exposure (**Figure 5C**). When crotonate or butyrate were added with fluconazole the expression of many *ERG* genes was reduced, as was the expression of *UPC2 and ECM22,* which encode transcription activators of *ERG* gene expression (**Figure 5C, Dataset S4**). Compared with SCFAs, TSA had less effect on *ERG* gene expression and no clear effect on *UPC2* and *ECM22* expression (**Figure 5C, Dataset S4**). Collectively, the transcriptional data indicates that SCFAs trigger reprogramming of acetyl-CoA metabolism in *C. albicans*, causing increased expression of β–oxidation genes and reduced expression of ergosterol biosynthesis genes, while TSA had a much milder effect on these metabolic pathways.

## DISCUSSION

Our study shows that peroxisome biogenesis genes and SCFAs, which are metabolised by peroxisomal β-oxidation affect fluconazole susceptibility in *C. albicans*. We propose that changes to peroxisomal metabolism, either by *pex* mutations or by supplementation of SCFAs, increases susceptibility to fluconazole by remodelling acetyl-coA metabolism and reducing ergosterol gene expression, thereby reducing the ability of cells to overcome fluconazole stress. A manuscript published while this study was in preparation supports this idea by showing that overexpression of *PEX8* reduces ergosterol gene expression and fluconazole tolerance (43).

Butyrate increases fluconazole susceptibility in *C. albicans,* and it also inhibits HDAC activity (26). Our results suggest that butyrate increases fluconazole susceptibility by HDAC-independent, as well as HDAC-dependent mechanisms. This conclusion is based on comparing two SCFAs, butyrate (a strong HDAC inhibitor) and crotonate (a weak HDAC inhibitor), as well as the strong HDAC inhibitor TSA, and finding that crotonate, the weakest HDAC inhibitor of the three compounds, synergised with fluconazole and strongly inhibited fluconazole tolerance to a greater degree than butyrate and TSA (**Figure 2**). Moreover, the RNAseq analyses found a shared transcriptional response to crotonate and butyrate, which included both upregulation of β-oxidation and peroxisome biogenesis genes and reduced expression of ergosterol biosynthesis genes.

Cells respond to fluconazole-induced inhibition of ergosterol biosynthesis by up-regulating *ERG* gene transcription (42); crotonate and butyrate reduce *ERG* gene expression levels (**Figure 5C**), providing an explanation for how these SCFAs synergise with fluconazole and how they eliminate tolerant growth (13, 26). TSA represses the expression of some *ERG* genes, such as *ERG1* and *ERG11* (**Figure 5C**) and (26), but the extent of repression was milder than the *ERG* gene repression mediated by SCFAs. For example, *ERG1* was more than 4-fold down-regulated in crotonate or butyrate in the presence of fluconazole relative to fluconazole alone, and less than 2-fold down-regulated by TSA+fluconazole relative to fluconazole alone (**Dataset S4**). Moreover, several *ERG* genes (e.g., *ERG251*, *ERG10*) did not undergo expression changes in TSA+fluconazole relative to fluconazole alone (**Figure 5C, Dataset S4** and interactive dataset <u>here</u>). Therefore, the SCFAs have a stronger effect on *ERG* gene repression than TSA, consistent with the idea that HDAC inhibition is not the only, and may not be the major mechanism by which SCFAs increase fluconazole susceptibility.

If and how increased β-oxidation in the presence of SCFAs is linked to reduced ergosterol biosynthesis remains to be fully understood. β-oxidation and ergosterol biosynthesis are metabolically linked, as acetyl-CoA produced by β-oxidation can serve as a precursor for ergosterol biosynthesis (**Figure 5A**). Therefore, increased β-oxidation in the presence of SCFAs would be expected to cause increased ergosterol biosynthesis. Indeed, in mice, increased β-oxidation caused increased cholesterol synthesis (44). However, we observed the opposite in *C. albicans*. While β-oxidation was upregulated in the presence of SCFAs, *ERG* gene expression was reduced. A possible explanation considers the need to balance ergosterol biosynthesis in yeast cells to avoid accumulation of toxic free sterols (45, 46). In other words, higher β-oxidation triggered by SCFAs may cause a feedback down-regulation of *ERG* gene expression, as a compensatory mechanism to balance the ergosterol biosynthesis pathway and prevent ergosterol overproduction. Under normal growth conditions (no drug), *C. albicans* grows well in the presence of SCFAs. However, when fluconazole inhibits ergosterol biosynthesis, the ability to upregulate *ERG* gene expression become critical. The transcriptomic data revealed that, in the presence of SCFAs *C. albicans* has a reduced ability to increase ergosterol gene expression that is required to overcome fluconazole. Therefore, we propose that balanced acetyl-CoA metabolism in the peroxisome is critical for *C. albicans* in achieving adaptive drug responses that ensure sufficient sterol biosynthesis to overcome fluconazole stress.

Our observation that butyrate increases fluconazole susceptibility in peroxisome biogenesis mutants (**Figure 2B**), suggests that SCFAs affect fluconazole responses not only through peroxisomal metabolism but also by additional mechanisms. These mechanisms are likely related to the ability of SCFAs to regulate gene expression via HDAC inhibition and by causing increased intracellular concentrations of acyl-CoAs that can modify histones and non-histone proteins. For example, upon supplementation of crotonate, increased histone crotonylation modulates gene expression and cellular responses (17, 19, 21, 24). Interestingly, a recent study reported that β-hydroxybutyrate synergises with fluconazole, possibly by promoting lysine-β-hydroxybutyrylation (47). Similarly, crotonate might increase fluconazole susceptibility by increasing lysine crotonylation, especially since the *C. albicans* proteome is heavily crotonylated (48). Given the strong effects of crotonate on fluconazole tolerance and synergism (**Figure 2**), it will be interesting to determine if lysine-crotonylation of histone or non-histone proteins regulates fluconazole responses.

In conclusion, this study adds to our growing understanding of the metabolic mechanisms that control antifungal drug susceptibility (2). Our experimental conditions used acidic pH to promote protonation of SCFAs and their uptake by fungal cells. These conditions mimic relevant host niches for *C. albicans*, including the vaginal tract. Therefore, SCFAs might have topical applications for increasing the efficacy of azole therapy in vulvovaginal infections.

## MATERIALS AND METHODS

### Strains and growth medium

The *C. albicans* strain SC5314 was used in most experiments. The *cph1Δ/*Δ *efg1Δ*/*Δ* mutant was obtained from Bernhard Hube (35) and construction of the *pex* mutants done in this study is described below. *Candida* was precultured from single colonies grown on YPD plates (2% peptone, 1% yeast extract, 2% glucose and 2% agar), inoculated into YPD liquid media and grown overnight at 30 °C at 200 rpm. Medium M199 was buffered with 3.5% HEPES at pH 5.4, including 80 µg/ml uridine. Crotonate and butyrate were used as sodium salts, and obtained by diluting concentrated crotonic or butyric acid with ddH_2_O before adjusting to pH 7.4 and filter sterilising. Stock Trichostatin A (TSA) was dissolved in DMSO to a concentration of 2 mg/ml. The synthetic complete (SC) medium was prepared as previously described (49). The medium was buffered to pH5.4 with 3.5% HEPES. The acetic, butyric, propionic and lactic acids were diluted in ddH2O adjusted to pH7.4, and filter-sterilised.

### Liquid filamentation assays

To observe hyphal filamentation in liquid media, cells were inoculated from stationary overnight cultures grown for 16 h at 30 °C, into M199 media at an OD_600_ of 0.1 and grown for 3 h at 37 °C. 1 ml of cells were then processed by centrifugation at 3000 rpm for 5 min. 500µl of supernatant was discarded. Cells were then centrifuged at 3000 rpm for 3 min and 200 µl of supernatant was discarded, twice. Cells were then centrifuged at 3000 rpm for 1 min and 50 µl of supernatant was discarded leaving cells concentrated in 50 µl of media. Cells were then mix pipetted and 3 µl of cells were mixed with 3 µl of mounting media solution and covered with a cover slip. Morphology was analysed at 40X and DIC images taken on Olympus microscope.

### Western blot analysis

Whole cell protein extraction and western blots were performed using our protocols described in (17, 23), with minor adjustments. To assay H3K9Ac and H3 for RNAseq conditions, *C. albicans* overnight cultures (YPD at 30 °C) were diluted to OD_600_ of 0.2 and harvested after 2 h in M199 medium at 37 °C and then treated for 30 min with or without 3 µg/ml fluconazole, 5 mM crotonate, 10 mM crotonate, 5 mM butyrate, 10 mM butyrate, 3 µg/ml TSA, 5 µg/ml TSA. All conditions were treated with equivalent amounts of DMSO. Samples were loaded on a 12% SDS-PAGE gel. Primary antibodies were H3K9Ac (Millipore, 07352) and H3 (Abcam, ab1791) at 1:5000 dilutions respectively. The HRP-conjugated anti-rabbit was the secondary antibody (1:20000 dilution).

### RNAseq

For the RNAseq experiment analyzing the response of *C. albicans* to SCFAs/HDAC inhibitors, overnight cultures of *C. albicans* (YPD at 30 °C) were diluted to OD_600_ of 0.3 into M199 (pH 5.4) and cells harvested after 2 h growth and following a 30 min treatment with 3 µg/ml fluconazole, 10 mM crotonate, 10 mM butyrate, 3 µg/ml TSA and combinations thereof. All conditions had equivalent concentrations of the solvent DMSO. RNA extraction was as described previously (17). Library preparation was by a modified version of the PAT-seq method (50). Briefly, 3’ barcoded cDNA was synthesised with Klenow Fragment (3’→5’ exo-) mediated extension of the 3’ end of polyA+ RNA, templated by an oligo containing a T12 3’ end to anneal to the poly(A) tail, followed by a 10bp UMI (Unique Molecular Identifier), 8bp barcode and truncated Illumina i7 sequence. cDNA was synthesised from end extended RNA with SuperScript III. Excess oligo was removed with thermolabile Exo I (NEB). The barcoded cDNA was pooled together, with 5 tubes processed in parallel with the equivalent of 1 microgram of RNA per tube, and RNA was removed by alkaline hydrolysis. Illumina i5 tags were added with a random primed 2^nd^ strand synthesis. Tagged 2^nd^ strand cDNA was size selected by urea-PAGE purification (to remove fragments smaller than 125 bp), and then amplified with barcoded Illumina i5 and i7 oligos. PCR products were gel purified to remove dimers. Samples were then checked for quality on Agilent Tapestation. Samples were sequenced on a NovaSeq 6000 with the SP Reagent Kit v1.5 (200 cycles), with a 202 bp (base-pair) (Read 1), 20 bp read (Read 2) and two 8 bp i5 and i7 index reads, with a 50% PhiX spike in for sequence diversity. UMI’s and barcodes were extracted from read 2 into the header of read 1 with UMI-tools: https://github.com/CGATOxford/UMI-tools. Reads were then demultiplexed with the bioinformatic tool demuxbyname.sh from the BBmap library and the samples were processed with the tail-tools package: https://github.com/Monash-RNA-Systems-Biology-Laboratory/tail-tools, alignment to the *C. albicans* reference genome SC5314 Assembly 21 were made with the STAR aligner (51).

Gene-wise read counts from tail tools were analysed in the interactive visualisation tool DEGUST. Genes with at least 10 reads and 5 CPMs in at least 3 samples were used for analysis. An apparent batch effect observed in dimension 2 of the MDS for Butr_R2 (**Figure 4A**) was caused by a slightly reduced RNA-seq library size and thus missing data for some low-expression genes. However, this did not impact the robustness of the statistical analysis based on genes that passed expression level filtering. To understand the impacts of SCFAs and TSA on the *C. albicans* transcriptome, the four groups of +crotonate, +butyrate, +TSA and control (DMSO) samples were selected together in the <u>DEGUST output</u>, analysed with ANOVA and the cut offs of >1.5 or 2 fold change and <0.05 false discovery rate (FDR) applied. Genes were then filtered by fold change in Excel to find those differentially expressed commonly or uniquely by the three treatments. The analysis is presented in **Dataset S2**. The same analysis was performed to understand how SCFAs and TSA impact on *C. albicans* in the presence of fluconazole, by using the +fluconazole condition as control and comparing it to +crotonate + fluconazole, +butyrate + fluconazole, +TSA+ fluconazole (**Dataset S3**). A third set of analyses was done by selecting five conditions (+DMSO, +fluconazole, +crotonate + fluconazole, +butyrate + fluconazole, +TSA+ fluconazole) and using either the DMSO or the fluconazole condition as control (**Dataset S4**). Pairwise comparisons of crotonate +/- fluconazole, butyrate +/- fluconazole and TSA +/- fluconazole were also performed and are presented in **Dataset S5**. Gene Ontology analysis was done using *Candida* Genome Database Term Finder and GraphPad Prism was used to construct heat maps. The RNAseq data has been deposited in GEO under accession number GSE303835.

### Disk diffusion assay

Overnight-grown cultures or single colonies were diluted and 100 µl plated on M199 or SDC plates +/- compounds using glass beads. A single 25 µg fluconazole disk (6mm diameter) was placed directly onto the middle of the plate using sterile forceps. Plates were incubated at 30 and 37 °C for 48 h and photographs taken on the Phenobooth+ (Singer Instruments). Quantification of disk diffusion assays was performed using *diskImageR* (52) or using J-AST, a JIPipe-reimplementation of diskImageR, where the data management is handled through the J-AST platform (https://asb.hki-jena.de/j-ast/) and JIPipe (53). *DiskImageR* is publicly available via CRAN (the Comprehensive R Archive Network). The Radius of Inhibition (RAD) values were calculated by distance in mm from the edge of the disk in relation to 20% inhibition (RAD_20_). The Fraction of Growth (FoG) (FoG_20_) within the zone of inhibition was determined using the area under the curve in slices from the disk edge to each RAD data point. This achieved growth is then compared to the potential growth.

### Fungal viability testing by colony forming units (CFUs)

Fungal viability in the presence of fluconazole, SCFAs and TSA was determined by counting colonies on agar plates at particular timepoints. Overnight-grown cultures were diluted to OD_600_ of 0.3 and grown in M199 at 37 °C for 2 h to an OD_600_ of 0.08-0.1. Following this, compounds were added at the indicated concentrations and samples were taken at various timepoints to optimise conditions for RNAseq. Serial dilutions were done to 1/2500, 1/5000 and 1/10000 to achieve reliable colony numbers suitable for counting. Plates were incubated on YPD plates at 30 °C for 2 days. A FIJI-based colony counting macro ‘ColonyCNTR’ (fun.qi – available at https://sites.google.com/monash.edu/fun-qi/home) was used to determine CFUs/ml.

### Minimal Inhibitory Concentration (MIC) and Checkerboard Assay

MICs were determined using the broth microdilution assay according to the CLSI guideline M27-A3, with minor adjustments. Single colonies of *C. albicans* were diluted to OD_600_ 0.1 in PBS and then diluted again to OD_600_ 0.01 into M199. Fluconazole concentrations ranged between 0.125 µg/ml to 16 µg/ml. Crotonate or butyrate concentrations ranged from 0.156 mM to 20 mM, or 0156 µg/ml to 20 µg/ml for TSA. In a 96 well plate, 50 µl of *C. albicans* inoculum was added to 50 µl of 2X drug concentration. A Tecan plate reader was used to measure the OD_600_ at the 24 h and 48 h timepoints. The MIC was determined by the concentration of fluconazole required to achieve a reduction in growth of 80% compared to the untreated control.

Antimicrobial synergy was determined through the checkerboard assay, concurrently in the same 96 well plate. Here, 25 µl *C. albicans* was added to 25 µl M199 and 25 µl of 4X solutions of both fluconazole and SCFAs or TSA. Fluconazole concentrations ranged between 0.25 µg/ml to 8 µg/ml in combination with 2.5 mM, 5 mM or 10 mM crotonate or butyrate, or 2.5 µg/ml, 5 µg/ml or 10 µg/ml of TSA. Growth was at 35 °C. A Tecan plate reader was used to measure the OD_600_ at the 24 h and 48 h timepoints. Combination effects were determined using the fractional inhibitory concentration index (FICI) method. Here, MIC_A_ and MIC_B_ reflect the MICs of the drugs alone and MIC_AC_ and MIC_BC_ correspond to the MICs of both drugs in combination, where FICI=(MIC_AC_/MIC_A_)+(MIC_BC_/MIC_B_). FICI values ≤0.5, 0.5 to <1, 1 to 4 or >4 are characterised as synergistic, additive, indifferent or antagonistic, respectively.

### Transposon library gDNA preparation and Illumina sequencing

The transposon library, previously described in (54) was plated on SDC-Ade medium supplemented with 10 µg/mL fluconazole and no drug (control). Plates were incubated at 30 °C for ∼3 days. Cells were harvested as previously described in (55), and used to inoculate 500mL of SDC-Ade liquid medium (± 10µg/mL FLC) to a final OD_600_ of 0.05. Cultures were incubated overnight at 30 °C at 220 rpm. Genomic DNA was extracted and prepared for Illumina sequencing as described previously (55) with minor modifications. Briefly, 2 µg of extracted genomic DNA was digested with 50 U of *NlaIII* restriction enzyme (New England Biolabs, USA). The self-ligation of digested fragments was done using 25 U of T4 DNA Ligase (New England Biolabs, USA). The Illumina library pools were done with inverse PCR methodology using two outward-facing primers, 3557-Illumina-RR and 3534-Illu_Tn_new_F (**Table S1**). The PCR amplification was performed in ten parallel 50µL reactions using PhireGreen Polymerase (Thermo Fisher Scientific, USA), which were subsequently pooled and purified using the NucleoSpin® Gel and PCR Clean-up kit according to manufacturer protocol (Macherey-Nagel, Germany). The resulting transposon pools were quantified with Agilent 4200 TapeStation system using HS D1000 ScreenTape. Illumina sequencing was performed at ZABAM NGS unit using NextSeq 2000 P3 XLEAP-SBS Reagent Kit 300 cycles (Tel-Aviv University). The raw sequencing data is available at NCBI under project PRJNA1477922 (http://www.ncbi.nlm.nih.gov/bioproject/1477922).

FASTQ files were processed as described previously in (54). Additionally, genes with low insertion coverage in control samples (≤10 insertion reads in 2 samples) were excluded prior normalization and differential analysis. The differential analysis was performed using the DESeq2 pipeline (56) estimating log_2_fold changes (log_2_FC), and p-values were assessed using the Wald test. Benjamini-Hochberg FDR correction was applied to control the false discovery rate.

### Construction of *C. albicans* deletion mutants

Deletion mutants were constructed in the *C. albicans* strain SC5314 using the transient CRISPR-Cas9 system described in (57), with modifications. To construct plasmid pJB523, we modified the CAS9-encoding plasmid pADH147 (57) by sequential removal of selection markers. The *NAT1* and *LEU2* markers were removed by sequential PCR amplifications, using primers MssI-487-FW + tCYC RV, and MssI-Cas9new-2976-FW + MssI-487-RV (**Table S1**), respectively, followed by a Gibson assembly.

To produce guide RNA fragments, the forward primer for amplification of the A fragment was replaced with primer pSNR52-F-2968, and the reverse primer for amplification of the B fragment was replaced by primer gRNAconserved-R. Genes were deleted using a ∼5kb repair fragment, containing the NAT resistance marker and a maltose-inducible FLP recombinase, was amplified from the plasmid pRB895-pSFS2A-mNeonGreen (58). Primers NAT-F and NAT-R carried 50-70 bp homology to the target ORF. *C. albicans* was transformed using the lithium acetate method and transformants were selected on YPD plates containing 200µg/mL NAT. After integration of the NAT-repair product into the genome, transformants were verified using PCR, the marker was removed at the FRT sites using FLP recombinase expression, induced by growing in YP + 2% maltose at 30 °C overnight, followed by replica-plating onto YPD and YPD plus 200µg/mL NAT plates to identify clones that lost the NAT-cassette. For each deletion, one correctly edited clone was selected for downstream analysis.

## ACKNOWLEDGMENTS

This work was supported by Future Fellowship awards from the Australian Research Council (ARC) to A.T. (FT190100733) and T.B. (FT180100049, CE230100001), and from the European Research Council (ERC) under the European Union’s Horizon 2020 research and innovation programme to J.B. (FungalTolerance 951475). We thank Bernhard Hube for the *cph1/efg1* mutant, Claudia Simm for advice on FICI calculations and Paul Harrison and David Powell from the Monash Genomics and Bioinformatic Platform for assistance with constructing RNAseq analyses and presentation with Degust. We also thank Nora Kawar, Dr. Anna Dukhovny for help in constructing mutants, and Dr. Shira Milo for help in data analysis.

## SUPPLEMENTAL DATA

**Figure S1.**
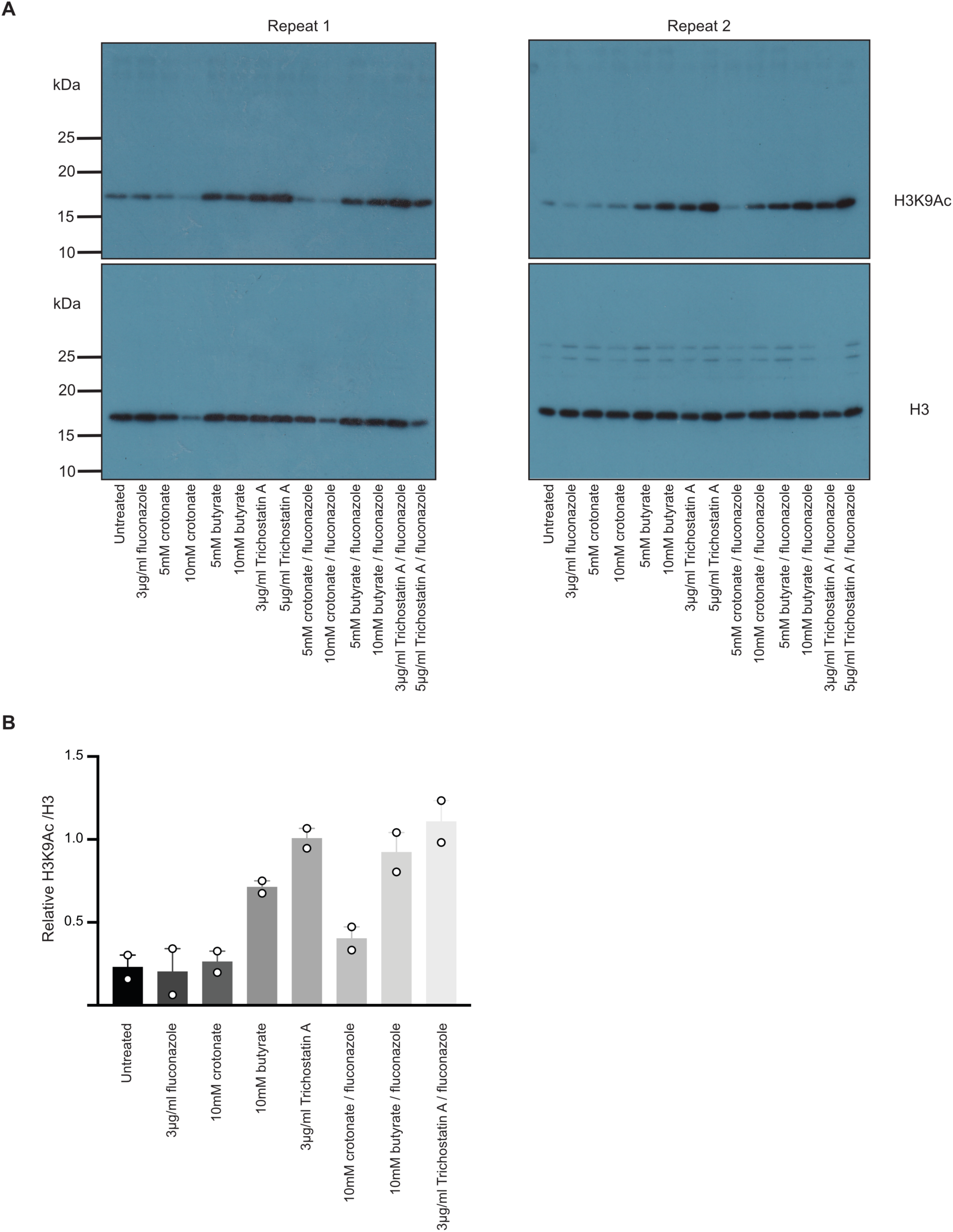
Western blot analyses of H9K9 acetylation upon treatment with crotonate, butyrate and TSA as well as fluconazole. **A.** Uncropped Western blots of two repeats analysing acetylation of histone 3 lysine 9 (H3K9Ac). Wild-type *C. albicans* SC5314 was grown for 2 h to log phase (OD 0.08-0.1) and then treated for 30 min with or without fluconazole in the presence or absence of crotonate, butyrate, and TSA at the indicated concentrations. Histone 3 (H3) was the loading control. **B.** Quantification of H3K9ac relative to total H3 from two independent experiments.

**Figure S2.**
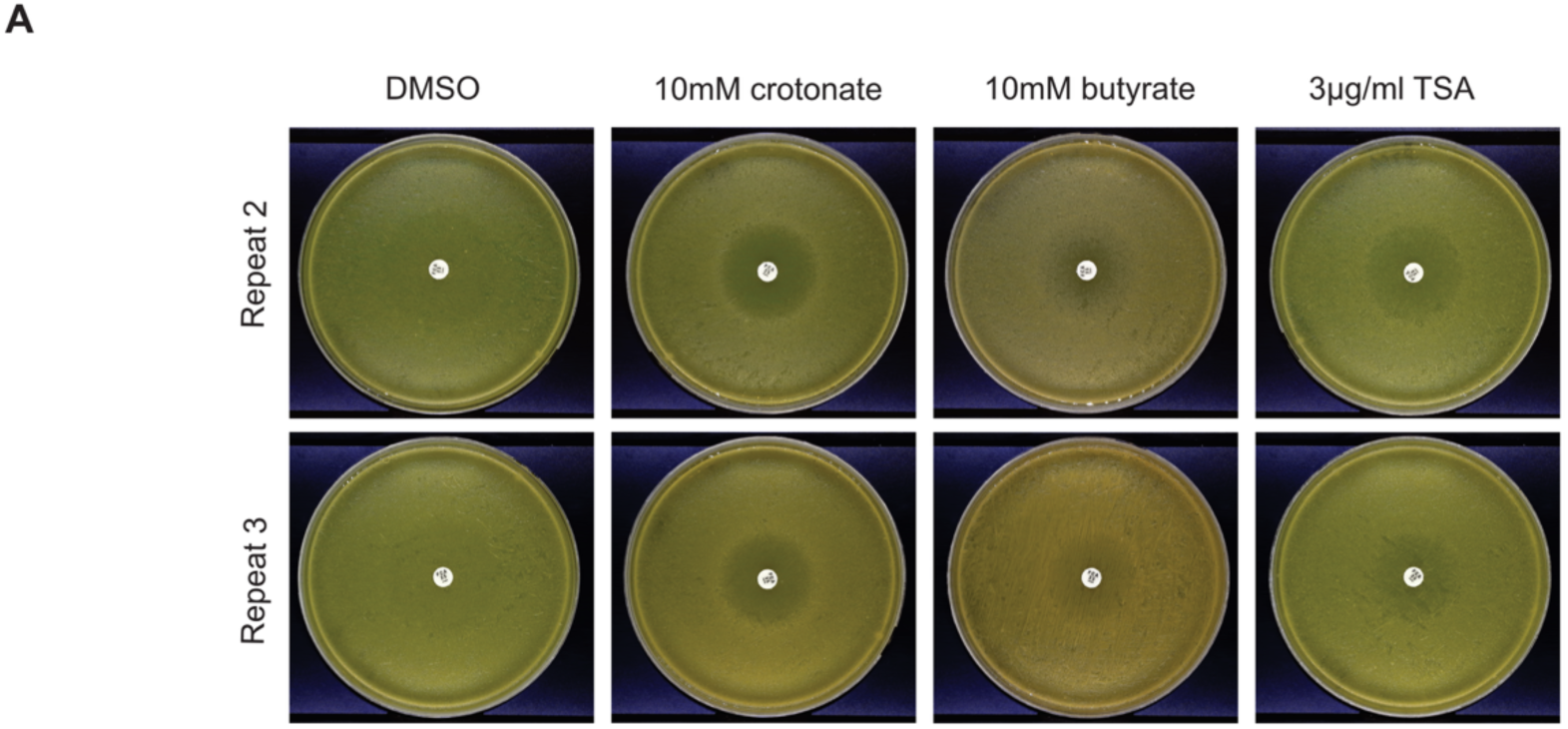
Additional experiments showing potentiation of fluconazole by crotonate, butyrate and TSA Fluconazole disk diffusion assays with or without 10 mM crotonate, 10 mM butyrate, 3 µg/ml TSA or equivalent DMSO vehicle control. The medium was M199, and the wild-type SC5314 strain of *C. albicans* was used. All plates contained equal amounts of DMSO. Plates were incubated at 37 °C and photographed after 48 h of incubation. Shown are images of three biological repeats. Repeat 1 is shown in Figure 2.

**Figure S3.**
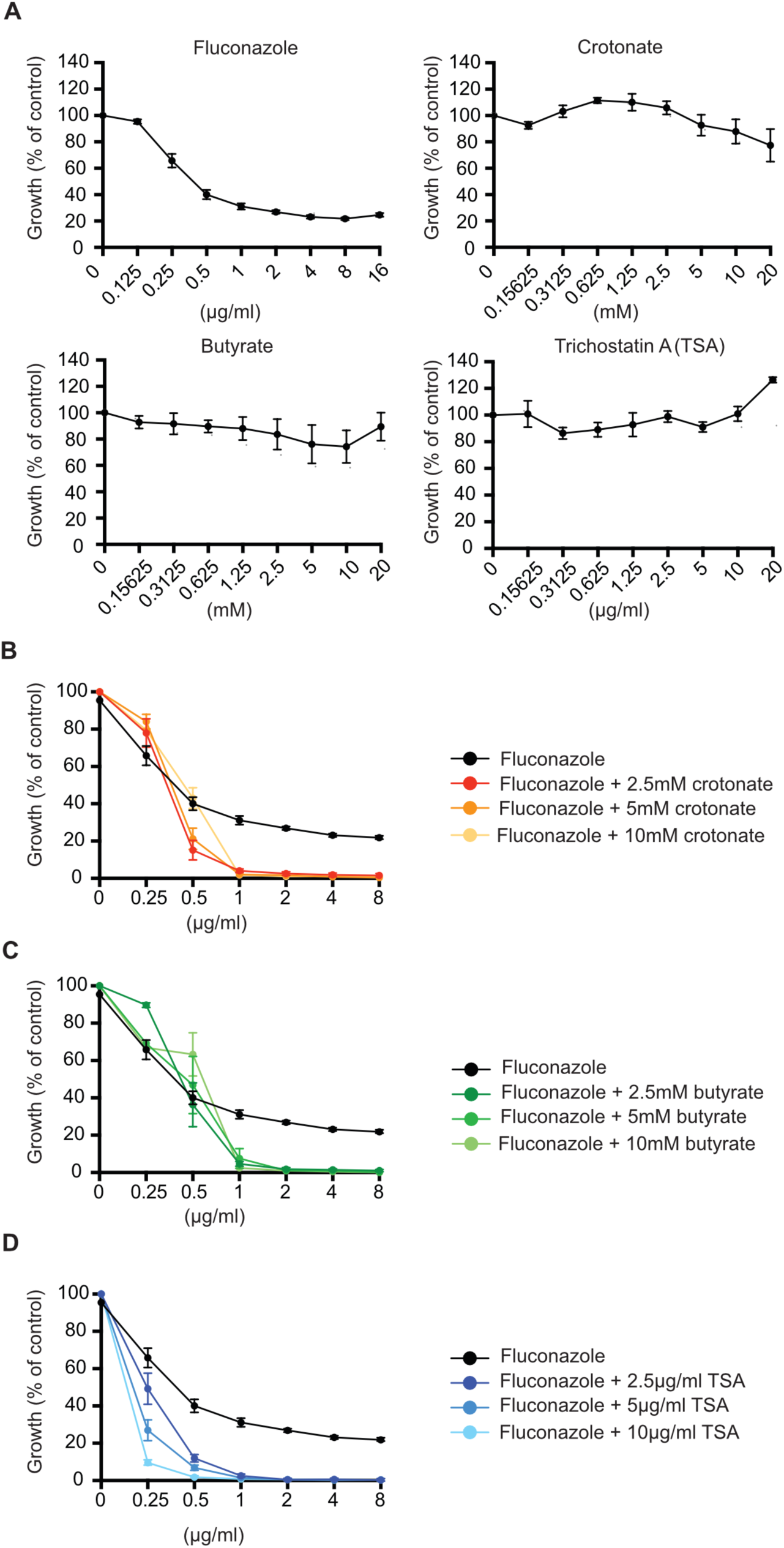
Growth data following combinatorial treatment with SCFAs, TSA and fluconazole Shown here is the growth data used to calculate the MICs, synergism, and construct the heat map in Figure 2. **A.** OD measurements for MIC assays with fluconazole, crotonate, butyrate and TSA at the indicated concentration. Wild type *C. albicans* (strain SC5314) was grown under a modified CLSI method at 35 °C in M199 and absorbance measured at OD_600nm_ at 24 h. **B.** As in A but for combinations of fluconazole with 2.5 mM, 5 mM or 10 mM crotonate. Graphs were plotted as a percentage of growth compared to the untreated control in M199 alone supplemented with equivalent volumes of DMSO. Data points are from three independent biological repeats. The fluconazole-only data is the same in panels B-D. These drug combinations were assayed together in the same experiments but are plotted separately for clarity. **C.** As in B but data is for co-treatment with butyrate. **D.** As in Band B but data is for co-treatment with TSA.

**Figure S4.**
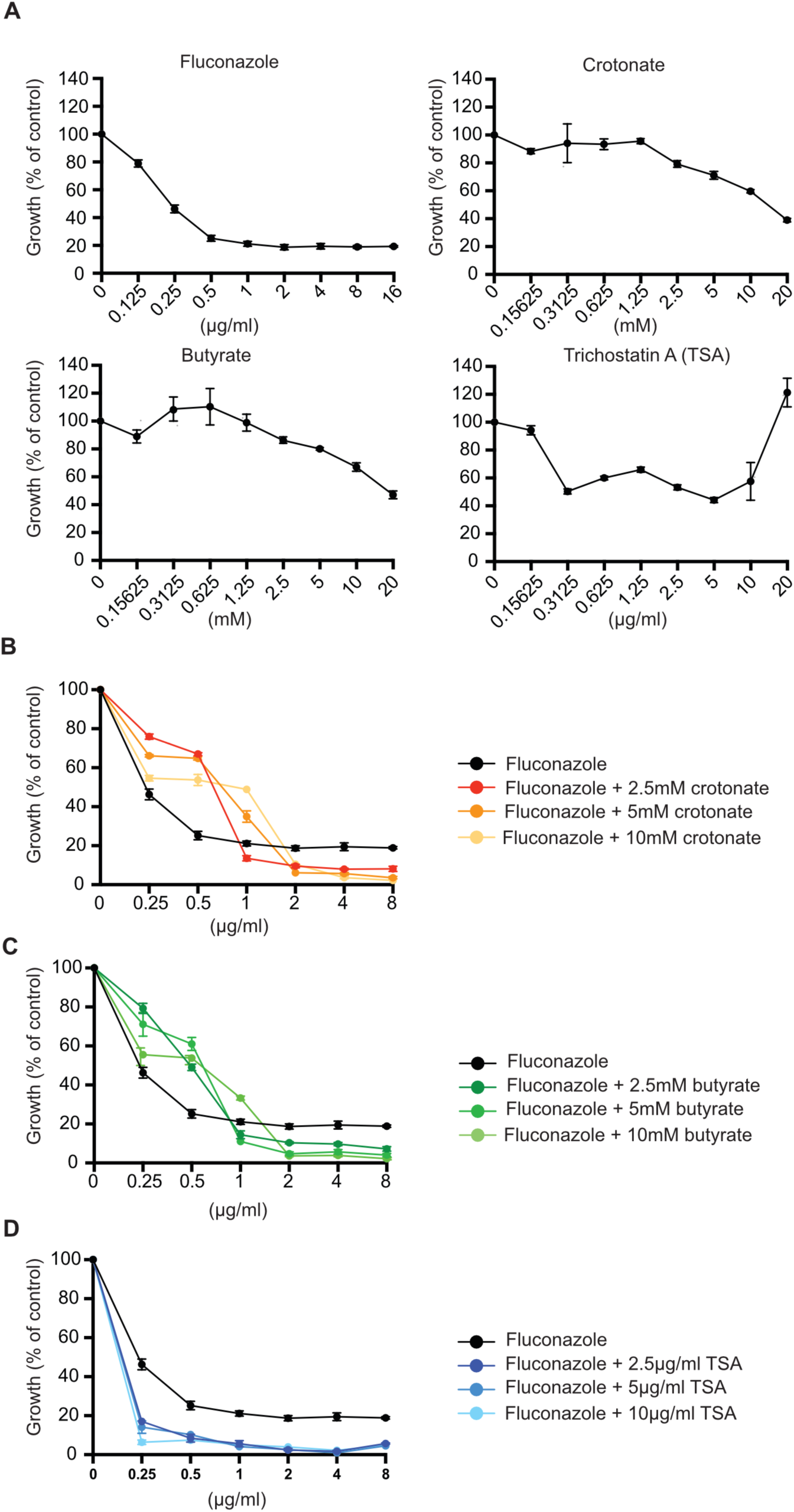
Growth data for combinations of the short chain fatty acids and TSA with fluconazole for the *cph1Δ/Δ efg1Δ/Δ* mutant Shown here is the growth data used to construct the heat map in **Figure 3**. **A.** Growth curves with fluconazole, crotonate, butyrate and TSA at the concentration indicated. The *cph1Δ/Δ efg1Δ* /*Δ* mutant cells were grown under a modified CLSI method at 35 °C in M199 and absorbance was measured at OD_600_ at 24 h. **E.** As in A but with fluconazole in combination with 2.5 mM, 5 mM or 10 mM crotonate. Absorbance for co-treatment was plotted against the absorbance produced in the fluconazole MIC assay by itself. Graphs were plotted as a percentage of growth compared to the untreated control in M199 supplemented with equivalent volumes of DMSO. Data points are from three independent biological repeats. The fluconazole-only data is the same in panels B-D. These drug combinations were assayed together in the same experiments but are plotted separately for clarity. **B.** As in B but data reflects co-treatment with butyrate. **C.** As in B but data reflects co-treatment with TSA.

**Figure S5.**
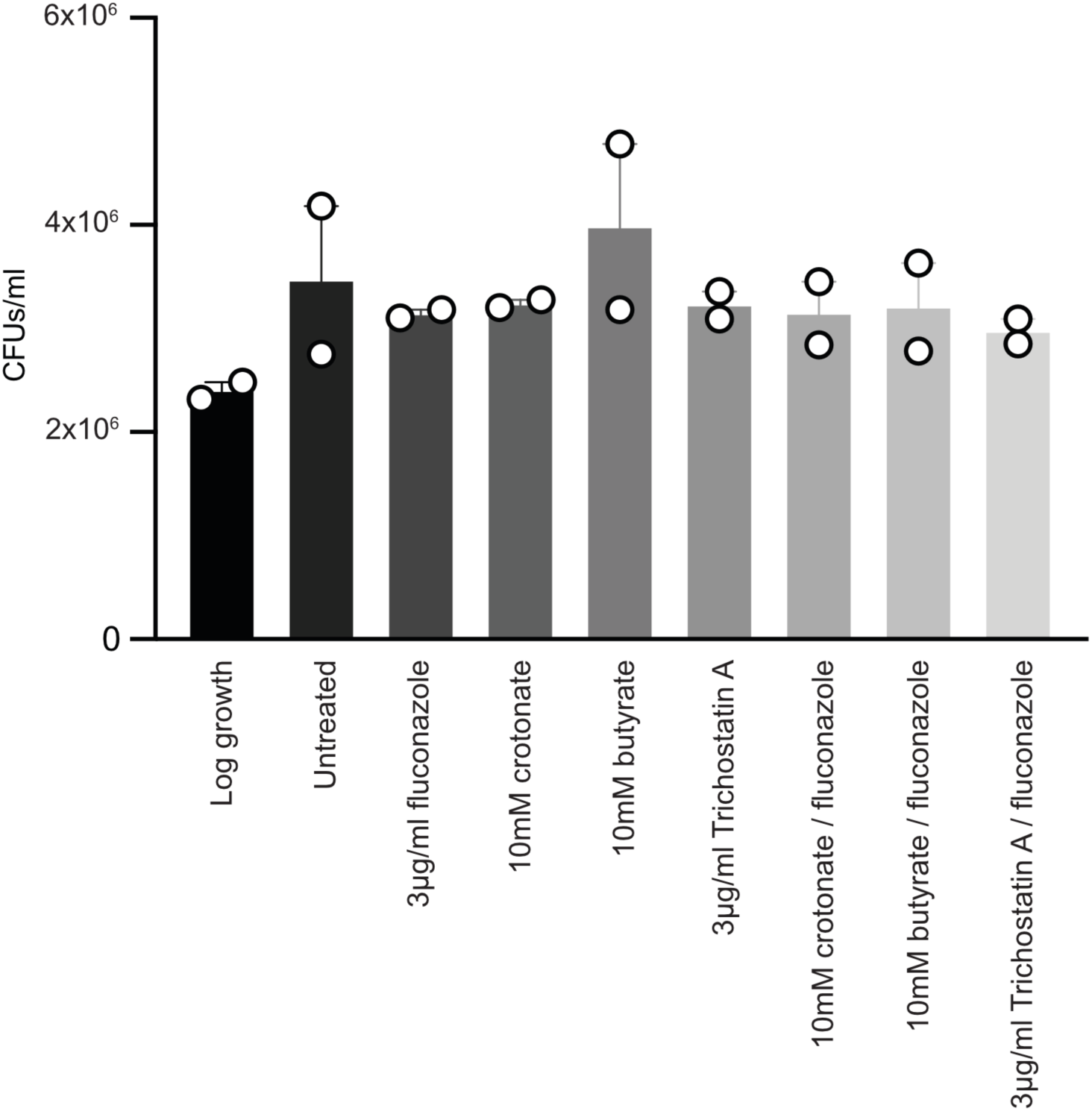
Assessing fungal growth under the conditions of the RNAseq experiments Colony forming units (CFU) analysis of *C. albicans* wild type SC5314 following 2 h growth to log phase (OD 0.08-0.1) and 30 min treatment with or without fluconazole in the presence or absence of 10 mM crotonate, 10 mM butyrate, 3 µg/ml trichostatin A. All conditions contained equivalent volumes of DMSO. CFUs were determined after plating onto YPD plates, and growth at 30 °C for 48 h. Data are from two independent experiments.

**Figure S6.**
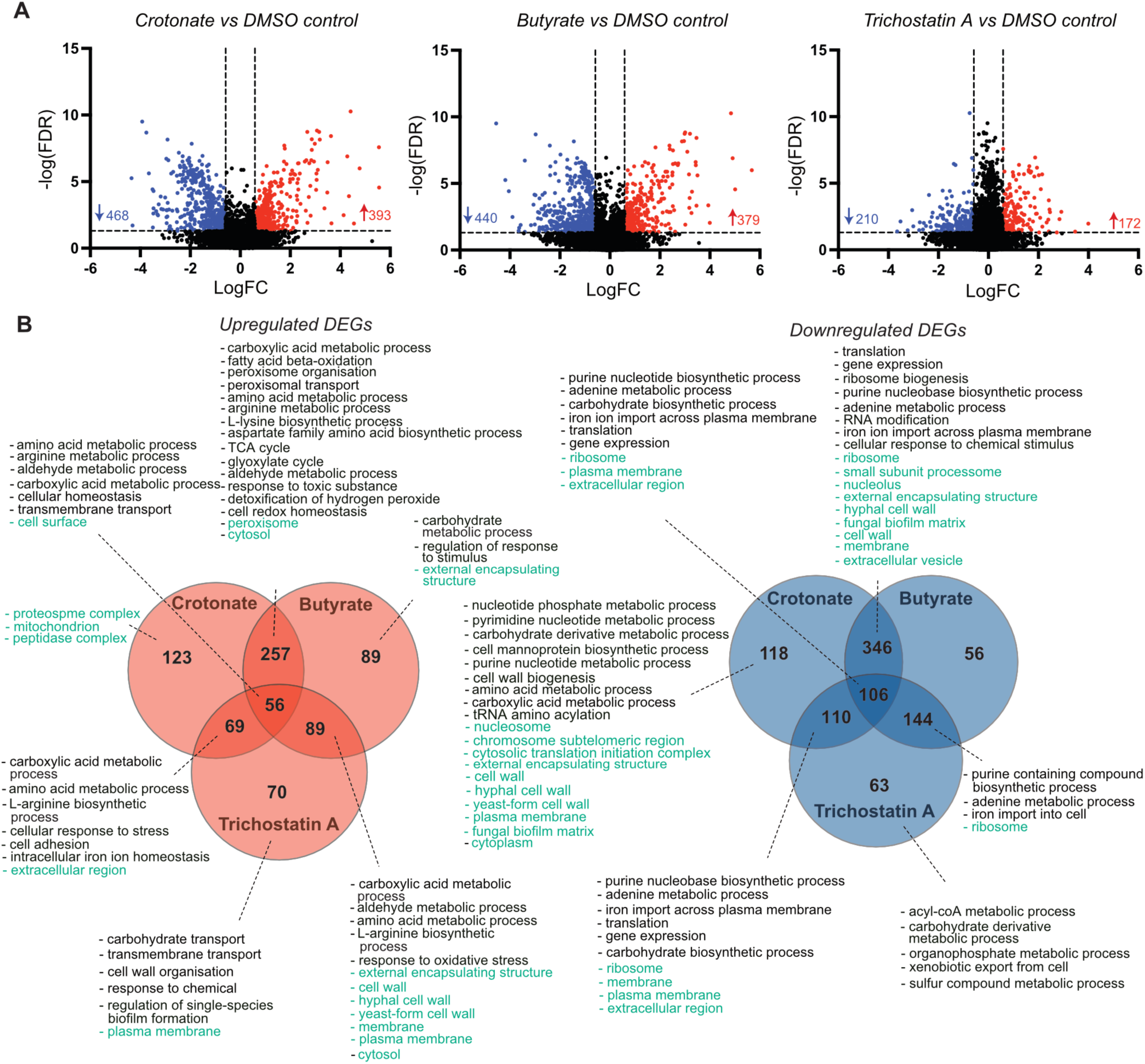
RNAseq data analyses of crotonate, butyrate and TSA treatment without fluconazole. **A.** Volcano plots of differentially expressed genes following treatment with crotonate, butyrate or TSA relative to DMSO control treatment (FDR 0.05, fold change of 1.5). **B.** Venn diagrams and functional terms for genes differentially expressed by butyrate, crotonate and TSA relative to the DMSO solvent control. The figure is based on Gene Ontology (GO) Term Finder analyses performed at the Candida Genome Database (p value of <0.1 used as a cut-off to assign hits). These analyses revealed many functionally overlapping GO terms (Process (black) and Component (green)) (see **Datasets S2** for full analyses). These functional terms were summarised here to avoid repetition and for simplicity. Up-regulated genes are shown in red; down-regulated genes are shown in blue.

---

**Table S1.** Primers used in this study

**Dataset S1. Transposon mutagenesis screen data**

**Dataset S2. RNAseq analyses: no fluconazole**

**Dataset S3: RNAseq analyses: in the presence of fluconazole**

**Dataset S4: RNAseq analyses: effects of SCFAs and TSA on acetyl-CoA and ergosterol metabolism**

**Dataset S5. Pair-wise comparison of the transcriptional changes caused by crotonate, butyrate and TSA in the presence and absence of fluconazole.**

